# Cloacimonadota metabolisms include adaptations for engineered environments that are reflected in the evolutionary history of the phylum

**DOI:** 10.1101/2021.10.08.463351

**Authors:** Lisa A. Johnson, Laura A. Hug

## Abstract

Phylum Cloacimonadota (previously Cloacimonetes, WWE1) is an understudied bacterial lineage frequently associated with engineered and wastewater systems. Cloacimonadota members were abundant and diverse in metagenomic datasets from a municipal landfill, prompting an examination of phylogenetic relationships, metabolic diversity, and pangenomic dynamics across the phylum, based on 22 publicly available genomes and 24 from landfill samples. Cloacimonadota formed two discrete clades, with one clade’s genomes principally deriving from engineered systems. A few more-divergent genomes were placed basal in the tree, and not associated with either clade. Metabolic reconstructions for metagenome-assembled genomes predict an anaerobic, acetogenic, and fermentative lifestyle for the majority of Cloacimonadota surveyed. Genomes from engineered ecosystems (first clade) encode a unique suite of genes not typically found in genomes from natural environments (second clade). These encoded functions include acetate kinase, the enzyme responsible for the formation of acetate from acetyl phosphate, and carbon utilization enzymes, suggesting different substrate degradation and energy generation strategies in these ecologically and phylogenetically distinct lineages.

**Originality/Significance Statement:** Cloacimonadota is a bacterial phylum that is under-described compared to its members’ prevalence in genome repositories. Cloacimonadota are frequently associated with engineered systems, including being identified as abundant and diverse in the municipal landfill site surveyed in this study. We reconstructed twenty-four landfill-associated Cloacimonadota metagenome assembled genomes (MAGs), more than doubling the number of publicly available Cloacimonadota genomes. We combined these MAGs with available reference genomes to predict major metabolic pathways and to describe the conserved features in the lifestyle of phylum Cloacimonadota. We found that Cloacimonadota have distinct evolutionary histories associated with engineered versus natural environments. Prior studies have evaluated metabolism from individual Cloacimonadota genomes – this work is the first to examine trait distribution across a more-complete representation of the phylum, including identification of genomic features and metabolic strategies that correlate to habitat of origin.

## Introduction

Cloacimonadota is an enigmatic bacterial phylum. Frequently observed in engineered systems like wastewater and sludge bioreactors, their metabolic diversity and adaptations to these man-made environments have not been examined. The first record of phylum Cloacimonadota was as unidentified 16S rRNA gene sequences from clone libraries derived from two unrelated samples: an anoxic flooded rice microcosm (Derakshani *et al*., 2001), and a reductive dechlorinating enrichment culture inoculated with material from a methanogenic reactor (Gu *et al*., 2004). Originally associated with the phylum Spirochaetota due to limited 16S rRNA gene reference sequences, Chouari *et al*., (2005) expanded the clade with sequences recovered from an anaerobic sludge digester to describe a well-supported monophyletic lineage distinct from Spirochaetota and other bacterial phyla (Chouari *et al*., 2005). The group was initially named candidate division WWE1, for the <u>w</u>aste<u>w</u>ater treatment plant in <u>E</u>vry, France from which the Chouari *et al*. 16S rRNA gene sequences were recovered, and as the first of two novel groups described in their study. The first, and only, complete genome for the phylum was sequenced from an anaerobic wastewater digestor (Pelletier *et al*., 2008). Provisionally classified as deriving from “*Candidatus* Cloacimonas acidaminovorans”, the genome was reconstructed using an iterative assembly process of genome extension from a metagenomic dataset (Pelletier *et al*., 2008). The genus name *Ca*. Cloacimonas derives from Latin terms translating to “a unit from the sewer”, with the species acidaminovorans translating to “amino acid eating”. In 2013, Rinke *et al*. reclassified the phylum from WWE1 to Cloacimonetes following expansion of the group with single-cell genomes from a brackish lake, deep sediment in a lagoon, and a terephthalate degrading bioreactors (Rinke *et al*., 2013). Cloacimonetes were a focus of Rinke and colleague’s work due to the phylum’s relative underrepresentation on the tree of life and the absence of a cultivated representative (Rinke *et al*., 2013), characteristics which hold true for the phylum today. Cloacimonetes were recently renamed to Cloacimonadota in the Genome Taxonomy Database (GTDB; http://gtdb.ecogenomic.org/) (Parks *et al*., 2018, 2020), a genome-based system attempting to standardized microbial taxonomy. The GTDB database currently contains 58 representative genomes for phylum Cloacimonadota, representing 23 species-level groups (Mendler *et al*., 2019). This level of available genomic diversity puts Cloacimonadota in the middle (48^th^ percentile) of the named bacterial phyla, further highlighting their comparative absence from the literature to date.

Cloacimonadota have frequently been identified from wastewater treatment plants (Westerholm *et al*., 2016; Calusinska *et al*., 2018; Jankowska *et al*., 2018; Theuerl *et al*., 2018; Shakeri Yekta *et al*., 2019) and biogas plants (Solli *et al*., 2014; Ahlert *et al*., 2016), often via 16S rRNA gene amplicon sequencing. Anaerobic digesters hosting Cloacimonadota span diverse wastewater substrates, including sugar beet molasses (Chojnacka *et al*., 2015), coffee processing (Botello Suárez *et al*., 2018), starch processing (Antwi *et al*., 2017; Qin *et al*., 2018), lipidic waste (Saha *et al*., 2019), cow manure/straw (Sun *et al*., 2015; Dong *et al*., 2019), municipal solid waste (Cardinali-Rezende *et al*., 2012; Akyol *et al*., 2019), sewage (Braz *et al*., 2019), and food waste (Moestedt *et al*., 2020). While Cloacimonadota are predominantly found in wastewater environments, they have also been found in the coal-bearing strata of the Cherokee basin (Kirk *et al*., 2015), an oilfield enrichment consortium (Toth and Gieg, 2018), and bodies of water including the meromictic Sakinaw Lake (Gies *et al*., 2014), a cold seep brine pool of the Red Sea (Zhang *et al*., 2016), the sulphidic zone in the Black Sea (Suominen *et al*., 2021), and the euxinic waters of Ursu Lake (Baricz *et al*., 2021).

The current understanding of Cloacimonadota metabolisms, lifestyles, and ecosystem roles largely derives from the “*Candidatus* Cloacimonas acidaminovorans” genome, where metabolism predictions indicate *Ca.* C. acidaminovorans is likely a syntrophic bacterium using fermentation of amino acids for carbon and energy (Pelletier *et al*., 2008). Another important study highlighting Cloacimonadota metabolism used enrichment cultures that included a member of Cloacimonadota, *Ca.* Syntrophosphaera thermopropionivorans (Dyksma and Gallert, 2019). Enrichment cultures were able to produce methane from propionate, and qPCR and 16S rRNA gene sequencing analyses validated the presence of *Ca.* S. thermopropionivorans from enrichments. Further, a draft genome of *Ca.* S. thermopropionivorans encoded some genes involved in propionate oxidation via methylmalonyl-CoA (Dyksma and Gallert, 2019). Other Cloacimonadota metagenome-assembled genomes (MAGs) derived from metagenomes of thermophilic biogas plants encoded the capacity to ferment amino acids with the production of acetate, H_2,_ and carbon dioxide (CO_2_) (Stolze *et al*., 2016), activities that were supported by metatranscriptomics for one of the Cloacimonadota MAGs (Stolze *et al*., 2018). Three Cloacimonadota MAGs from deep terrestrial subsurface sediments contain [FeFe] hydrogenases, suggesting members of this group have potential roles in H_2_ cycling (Hernsdorf *et al*., 2017). Other studies indicated that members of the Cloacimonadota have cellulolytic potential (Limam *et al*., 2014; Xia *et al*., 2018), and quantitative DNA stable isotope probe incubations along with a MAG recovered from the sulphidic zone of the Black sea highlighted that one Cloacimonadota member is a generalist capable of using different organic matter substrates (Suominen *et al*., 2021).

Consistent with strong representation in engineered waste environments, Cloacimonadota have also been identified at municipal landfills. Cloacimonadota have been observed at mature landfill sites where microbial communities are involved in humification (Liu *et al*., 2011; Remmas *et al*., 2017), and as part of landfill leachate microcosms enriched for lignocellulose degradation (Ransom-Jones *et al*., 2017). However, some investigations of bacterial diversity at municipal landfills did not detect Cloacimonadota (Song *et al*., 2015; Stamps *et al*., 2016), which could be due to the heterogeneous nature of municipal solid waste (MSW) at landfills. The drivers for Cloacimonadota presence or absence within a municipal landfill are not known.

Our study site is an active municipal landfill in Southern Ontario. The landfill is instrumented with several hundred wells for monitoring leachate chemical composition. Leachate is the liquid component of landfill waste, derived from microbial degradation of organic wastes, rainwater inflow, and groundwater infiltration, where leachate typically contains many organic and inorganic compounds, with the composition varying with the make-up of deposited waste (Renou *et al*., 2008). These compounds can have adverse impacts on surrounding environments if leachate is not properly contained or treated. Alongside the formation of leachate, microbially-mediated decomposition of MSW results in gas formation, including the greenhouse gases CO_2_ and methane (CH_4_) (Hilger and Barlaz, 2006). To mediate the negative impact of gas emissions, many modern landfills, including our study site, are instrumented with gas capture systems that redirect emitted methane to biogas generation plants, subsequently converting methane to electricity (Spokas *et al*., 2006). At the Southern Ontario landfill, the gas capture system experiences periodic fouling by overgrowth of a thick biofilm (personal communication, landfill engineers). Biofouling blocks the methane-capture infrastructure, consequently reducing methane gas capture and sustainable energy production. The infrastructure at the landfill site uses leachate wells to monitor leachate, composite leachate cisterns to collect leachate formed across the site prior to transport to a wastewater treatment facility, and a gas capture system to collect methane and other gases for bioenergy conversion. Diverse and abundant Cloacimonadota have been detected in leachate wells (LW), a composite leachate cistern (CLC), and a biofilm (BF) within the gas capture system across the site (Sauk and Hug, 2021).

Here we expand the genomic and metabolic diversity of the understudied phylum Cloacimonadota using MAGs derived from landfill metagenomes. We examine the metabolic capacities and roles for this lineage within engineered environments. In a pangenome comparison of all currently available Cloacimonadota genomes, we assess traits specific to organisms found in anthropogenically-impacted or pristine environments. Our results delineate the ecosystem services the members of this phylum contribute to an environment and clarify why these organisms are so frequently prevalent in bioreactors and other engineered, anaerobic environments.

## Results and discussion

### Characteristics of Cloacimonadota genomes

The genome quality of all landfill-derived and publicly available Cloacimonadota genomes was evaluated using CheckM (Table S1) (Parks *et al*., 2015). Of the 29 Cloacimonadota MAGS from the landfill datasets, 24 were of sufficient quality (>70% completion, <10% contamination). From public databases, 22 of 40 Cloacimonadota genomes and MAGS were retained for further analyses. The highest quality genome is, unsurprisingly, from the sole sequenced isolated, “*Candidatus* Cloacimonas acidaminovorans” strain Evry, with a completion of 100% and predicted contamination of 1.1% (Pelletier *et al*., 2008). Within the set of Cloacimonadota genomes kept for further analyses, the number of scaffolds ranges from 1 to 443, with a median of 135 (Table S1). The average genome size was 2.4 Mbp, with an average of 2,028 genes per genome. Generally, the number of predicted genes increases linearly with genome size, irrespective of which environment the genome originated from (Figure 1). The largest genome belongs to *Candidatus* Cloacimonetes bacterium NORP72 from marine subsurface, with a size of 4.6 Mbp (Tully *et al*., 2018), and the smallest genome belongs to *Candidatus* Cloacimonetes bin NIOZ-UU4 from an epipelagic marine environment, with a size of 1.4 Mbp (Villanueva *et al*., 2021).

**Figure 1.**
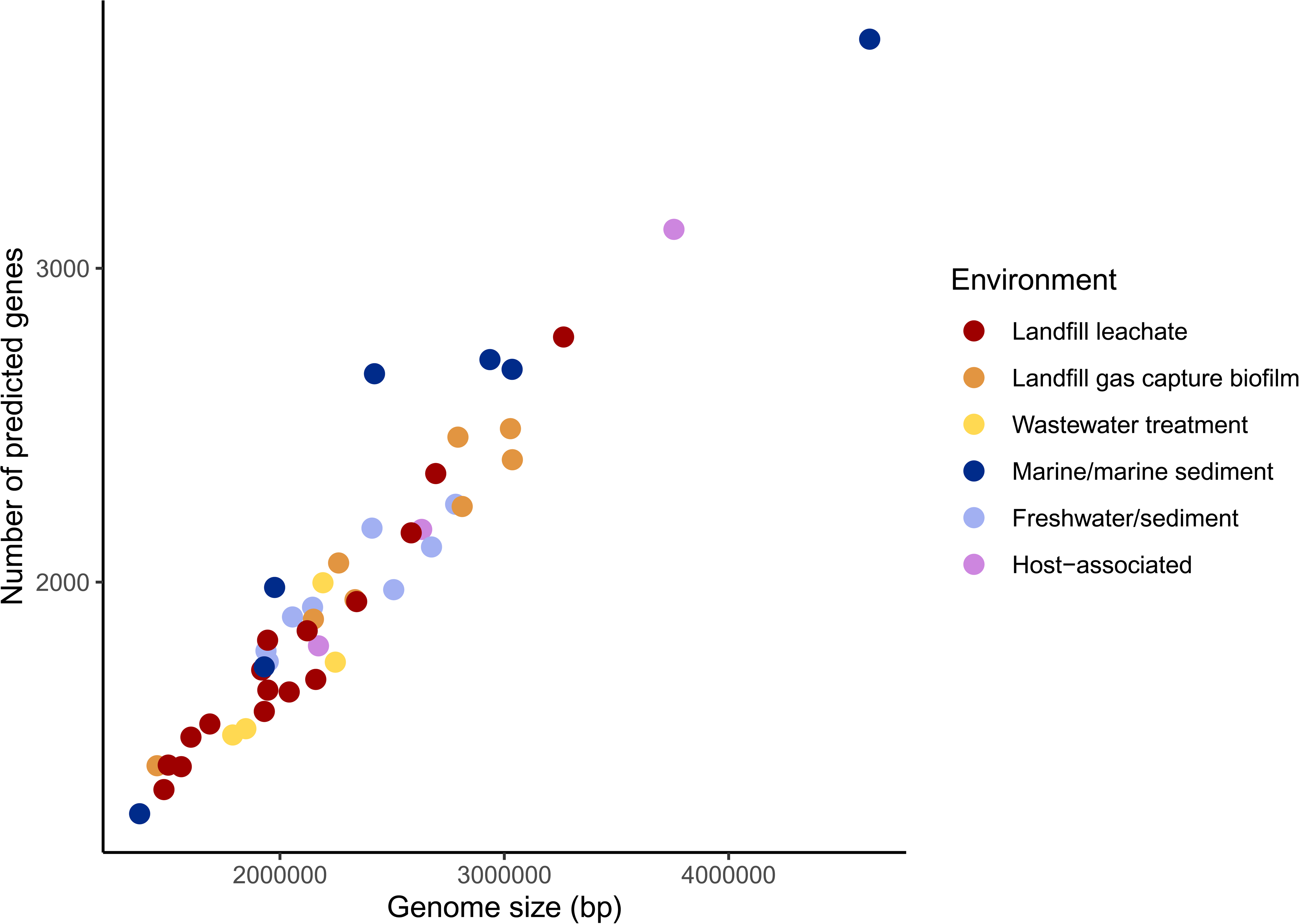
Number of predicted genes compared to genome size for high quality Cloacimonadota genomes (n = 46). Each Cloacimonadota genome is represented by a dot, colored by the environment of origin.

### Environmental distribution of Cloacimonadota

Cloacimonadota were detected at different locations with the waste management system in Southern Ontario (Table S1). At the landfill site, Cloacimonadota MAGs were represented in metagenomes from three leachate wells (LW1, n = 2; LW2, n = 8; LW3, n = 1), the composite leachate cistern at both time points (CLC_T1, n = 3; CLC_T2, n = 7), and the biofilm sample (BF1, n = 8) (Table S1). Cloacimonadota were not represented in the MAGs from the groundwater well metagenome at the landfill site. The 24 MAGs passing the CheckM quality metrics were analyzed further. The proportion of reads from the total assembled dataset attributed to each MAG provides an estimate of relative abundance across landfill sites. Across the landfill sites, Cloacimonadota MAGs accounted for 0.06% to 4.72% of the assembled metagenome, with an average of 0.44%, making them numerous but not notably abundant members of the landfill microbial communities. The biofilm MAG, BF1 Bin 72, had the highest proportional abundance within a sample by far, with 4.72% of the reads included in the assembled metagenome mapping to this genome. The next highest proportionally abundant MAG, BF1 Bin 89, contained 0.71% of the assembled metagenome. A total of 22 moderate quality Cloacimonadota genomes were publicly available, generated from diverse locations and ecosystems (Figure 2; Table S1). The addition of 24 MAGs from the Southern Ontario landfill expands the available genomic representation of Cloacimonadota by 109%, a substantial increase for this radiation.

**Figure 2.**
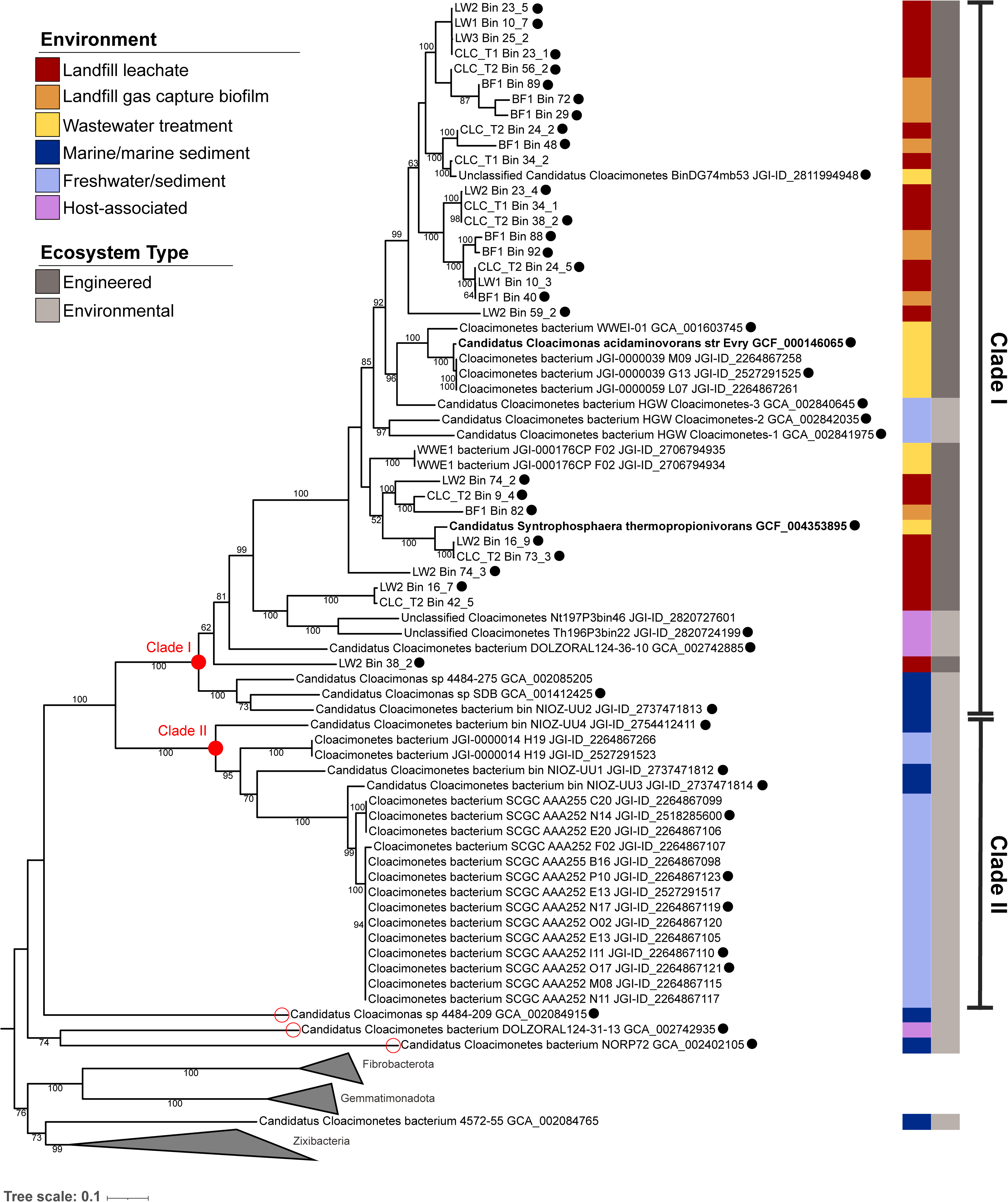
Phylogenetic placement of Cloacimonadota genomes. A maximum likelihood tree was inferred based on a concatenated alignment of 16 ribosomal proteins. Gemmatimonadota, Fibrobacterota, and Zixibacteria genomes were included as outgroups. The outside greyscale bar indicates the type of ecosystem from which the genome was obtained, and the inner colored bars further discern the specific environment of origin. Black circles indicate genomes that were of sufficient quality (>70% completion, and <10% contamination), and were included in further metabolic analyses (n = 21, with landfill MAGs n = 46). The two bolded genome names denote two genomes which have been previously investigated regarded metabolic function (Pelletier *et al*., 2008; Dyksma and Gallert, 2019). The nodes that anchor Clade I and Clade II are highlighted with red circles. Three genomes did not fall into either clade, and are indicated with red circles on the tips beside the genome name.

Phylogenetic analyses based on 16 concatenated ribosomal proteins identified all genomes (regardless of quality metrics, n= 68) as belonging to the monophyletic phylum Cloacimonadota. The phylogenetic tree displays Cloacimonadota genomes falling into two clades differentiated by ecosystem type, where one clade contains genomes derived predominantly from engineered ecosystems (Clade I), and the other clade contains genomes derived from natural, though not necessarily unimpacted, environmental ecosystems (Clade II) (Figure 2). Clade I is a monophyletic lineage (bootstrap = 100, total n = 46, passing quality n = 35 (black circles)), composed primarily of genomes isolated from engineered ecosystems (37/46) and includes all landfill-derived MAGs. Other genomes within this engineered clade were reconstructed from wastewater treatment plants (n = 6) of varying types: municipal (Pelletier *et al*., 2008), terephthalate-degrading (Rinke *et al*., 2013), cellulose-degrading (unpublished), and solid organic waste (unpublished). Two genomes were from a biogas plant reactor metagenome (unpublished). Clade I also contained six genomes from natural ecosystems, and three that are host-associated (Figure 2). Three groundwater genomes were derived from a sediment ecosystem dedicated to radioactive waste disposal research (Hernsdorf *et al*., 2017). While this groundwater system is classed as a natural environment, the sampled location is anthropogenically affected. Another three genomes in Clade I were isolated from animal hosts: two from different termite gut samples (unpublished) and the last from a dolphin’s oral cavity (Dudek *et al*., 2017). The last three genomes encompass a small subgroup at the base of Clade I (bootstrap = 100), and are *Candidatus* Cloacimonas sp. 4484_275, *Candidatus* Cloacimonas sp. SDB, and *Candidatus* Cloacimonetes bin NIOZ-UU2, which originated from deep sea hydrothermal vent sediments (Dombrowski *et al*., 2018), a methanogenic culture enriched from contaminated sediments (Callaghan *et al*., 2010), and the Black sea at a depth of 70 m (Villanueva *et al*., 2021), respectively. Placed basal within Clade I, these three environmental genomes were of specific interest when examining the evolution or acquisition of traits affiliated with adaptation to engineered environments.

Clade II is a monophyletic group of 19 genomes from environmental ecosystems (Figure 2; bootstrap = 100). Of those genomes, 14 are single-cell genomes from Sakinaw Lake, collected from a depth of 120 m, a location that is brackish, devoid of oxygen, and has high levels of hydrogen sulfide (1000 µM) (Rinke *et al*., 2013). From the same study, Clade II also contains two genomes from lagoon sediment that is similarly devoid of oxygen and with high levels of hydrogen sulfide (Rinke *et al*., 2013). The remaining three genomes from this clade originate from the water column from deep sea dives in the Black Sea (Villanueva *et al*., 2021).

Beyond these two clades, there are three genomes that were more distantly related to other Cloacimonadota genomes than to each other, at the base of the Cloacimonadota radiation though still clearly associated with the phylum (Figure 2). *Candidatus* Cloacimonetes bacterium DOLZORAL124_31_13 originated from a dolphin’s oral cavity (Dudek *et al*., 2017), *Candidatus* Cloacimonetes bacterium NORP72 from an oxic subseafloor aquifer (Tully *et al*., 2018), and *Candidatus* Cloacimonas sp. 4484_209 from marine deep-sea hydrothermal vent sediments (Dombrowski *et al*., 2017). For metabolic comparisons, these three genomes were considered separate from both Clade I and Clade II. One genome did not fall within the Cloacimonadota monophyletic lineage: *Candidatus* Cloacimonetes bacterium 4572_55 (Dombrowski *et al*., 2017) branched as a sibling to the Zixibacteria, within the Zixibacteria/Fibrobacterota/Gemmatimonadota outgroup. This calls this genome’s designation as a Cloacimonadota into question, and it was excluded from further metabolic analyses.

### Metabolic potential of Cloacimonadota

The Cloacimonadota landfill MAGs and reference genomes passing quality standards (>70% completion, <10% contamination; n = 46) were analyzed with the KEGG Automatic Annotation Server (KAAS) (Table S2) (Moriya *et al*., 2007), and the program called Distilled and Refined Annotation of Metabolism (DRAM) (Table S3 and Figure S1) (Shaffer *et al*., 2020), each of which provides an overview of the metabolic functions encoded by each genome. More detailed examinations for specific functions of interest were carried out through manual curation.

### General lifestyle

Cloacimonadota are predicted to be anaerobic, gram-negative bacteria (Pelletier *et al*., 2008). Gram-negative bacteria have two membranes: an inner membrane surrounding the cytoplasm and an outer membrane protecting the cell from the environment that is made from lipopolysaccharide (LPS) molecules (Bertani and Ruiz, 2018). Based on KEGG annotations, all 46 genomes encoded partial pathways involved in lipopolysaccharide biosynthesis, such as the Kdo_2_-lipid A biosynthesis pathway (Raetz pathway), indicating these organisms are likely gram-negative. It was previously described that members of phylum Cloacimonadota encode genes involved in generating archaeal-like membrane lipids, specifically geranylgeranylglyceryl phosphate (GGGP) and (*S*)-2,3-di-O-geranylgeranylglyceryl phosphate (DGGGP), which may allow for the generation of mixed archaeal/bacterial membranes (Villanueva *et al*., 2021). GGGP (K17104) and DGGGP (K17105) were encoded in 24% (11/46) and 30% (14/46) of Cloacimonadota genomes, with 24% of genomes encoding both genes. The genomes encoding GGGP and DGGGP in Clade I include LW2 Bin 16_7, Unclassified Cloacimonetes Th196P3bin22, *Candidatus* Cloacimonetes bacterium DOLZORAL124_36_10, LW2 Bin 38_2, *Candidatus* Cloacimonas sp. SDB, and *Candidatus* Cloacimonetes bacterium bin NIOZ-UU2, all of which are deep-branching within Clade I, suggesting this trait was lost in later-diverging lineages. Another important cellular component is peptidoglycan, which provides structural support for maintaining cell shape, and the *mur* enzymes involved in peptidoglycan biosynthesis are partially represented in all Cloacimonadota genomes. The complete pathway was only detected in 6% of genomes from the engineered Clade I (2/35 genomes), 63% of the environmental Clade II genomes (5/8), and 100% of the remaining, basal Cloacimonadota genomes (3/3)

Cloacimonadota genomes encoded genes for type IV pili, including genes involved in the components for the major pilin, assembly ATPase, retraction ATPase, platform protein, and alignment protein (Craig *et al*., 2019). Additionally, many genes involved in flagellar systems were encoded by a select group of five Cloacimonadota genomes, including LW2 Bin 16_7, Unclassified Cloacimonetes Th196P3bin22, *Candidatus* Cloacimonetes bacterium DOLZORAL124_36_10, *Candidatus* Cloacimonetes bacterium DOLZORAL124_31_13, and *Candidatus* Cloacimonetes bacterium NORP72. The flagellar genes encoded in these five genomes are consistent with genes found to be ‘core’ in diverse and flagellated bacteria from 11 different phyla (Liu and Ochman, 2007).

The most common defenses against reactive oxygen species (ROS) encoded in Cloacimonadota genomes are thioredoxin reductase (K00384), and thioredoxin 1 (K03671), which were each present in 89% (41/46) of genomes. Alkyl hydroperoxide reductase (K03386) was encoded in 59% (27/46) of Cloacimonadota genomes. Thiol peroxidase (K11065) was detected in 74% (26/35) of Cloacimonadota genomes from Clade I, and none of the genomes from Clade II. Similarly, glutathione peroxidase (K00432) was encoded in 40% (14/35) of genomes from Clade I, and none from Clade II. Other ROS defenses were detected sporadically across Cloacimonadota genomes or were not detected at all (Figure 3). Based on the KEGG annotations, superoxide reductase (K05919) and rubrerythrin (K19824) were not detected in Cloacimonadota genomes though these genes were encoded in *Ca.* C. acidaminovorans and *Ca.* S. thermopropionivorans (Dyksma and Gallert, 2019). Therefore, a manual BLAST search was conducted using superoxide reductase (WP_133200558.1) and rubrerythrin (WP_133200557.1) genes annotated in the *Ca.* S. thermopropionivorans genome, where the genes were detected in 83% (38/46) and 89% (41/46) of the genomes, respectively. The distribution of genes in Cloacimonadota genomes is generally consistent with a previous broad search of ROS defenses across bacterial phyla (Johnson and Hug, 2019), with the exception of rubrerythrin, which was not detected in the broader search, but was identified here under a targeted search. It is possible there was enough sequence dissimilarity for superoxide reductase and rubrerythrin to be missed with the automated KEGG annotation, though both were detected with manual BLAST searches. Collectively, for both Cloacimonadota clades, management of damaging free radicals appears limited, likely constraining these organisms to anoxic environments.

**Figure 3.**
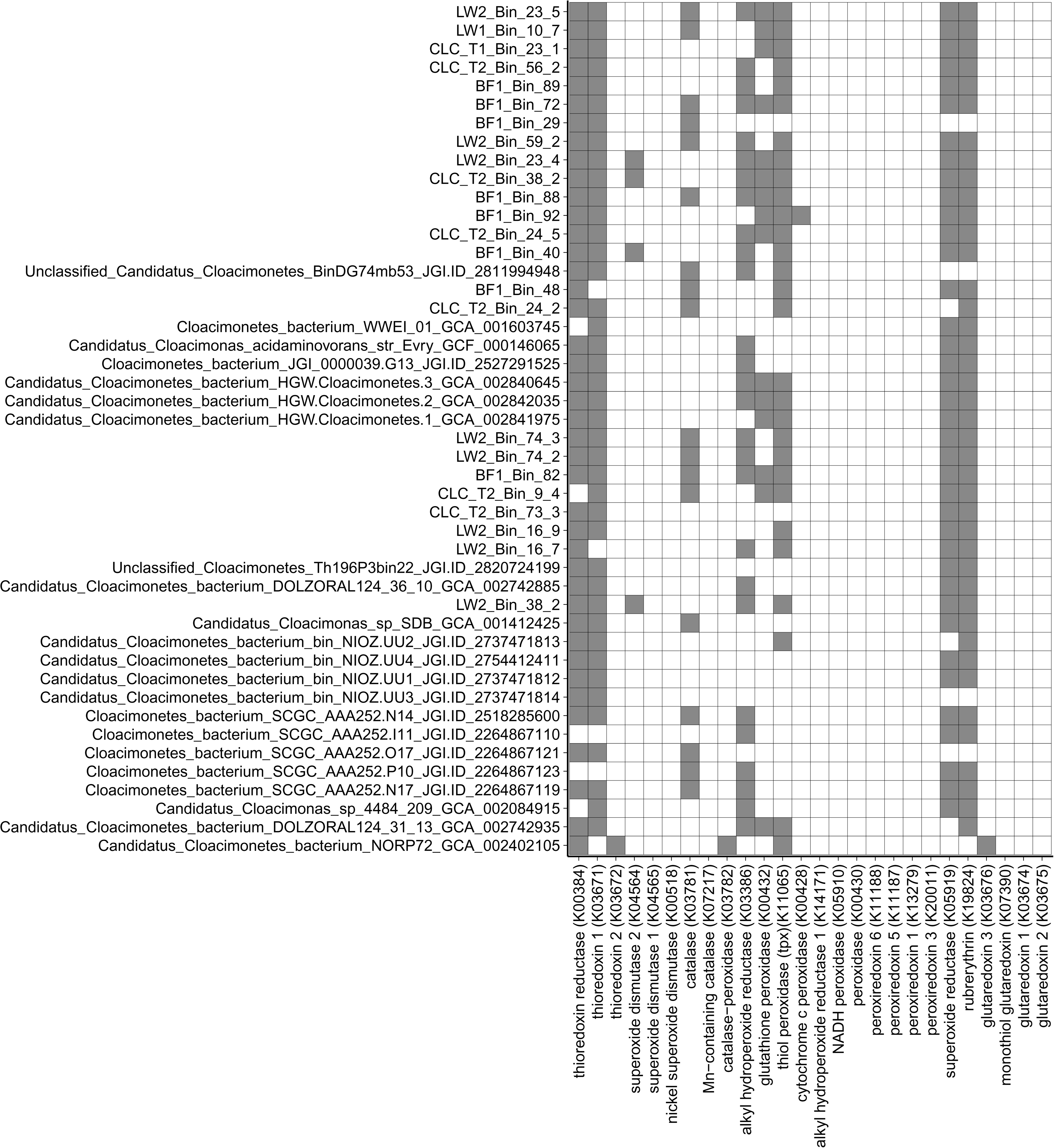
Reactive oxygen species (ROS) defense gene presence or absence in Cloacimonadota genomes. Dark grey boxes denote ROS defense genes that were encoded, and white indicates ROS genes that were not detected from the KEGG annotations or manual screens for each genome.

### Carbon metabolism

The Cloacimonadota genomes generally contain a full glycolysis pathway, with the notable exception of pyruvate kinase, which catalyzes the formation of pyruvate. The other nine enzymes of the glycolysis pathway were found in 72% to 98% of genomes (33 to 45/46), whereas pyruvate kinase (EC: 2.7.1.40) was only identified in 26% of the genomes (12/46), and not associated with any specific phylogenetic radiation. To ensure pyruvate kinase was not missed during automated annotation or due to partial hits, additional BLAST searches of Cloacimonadota genomes for pyruvate kinase were conducted, which did not locate any additional genes. While 74% of Cloacimonadota genomes were missing pyruvate kinase, pyruvate orthophosphate dikinase (EC: 2.7.9.1), which catalyzes a reversible reaction between pyruvate and phosphoenolpyruvate (PEP), was present in 91% of the genomes (42/46). Pyruvate orthophosphate dikinase has been shown to mediate pyruvate flux in *Clostridium thermocellum* (Olson *et al*., 2017), and we predict a similar role in the Cloacimonadota. The presence of either pyruvate kinase or pyruvate orthophosphate dikinase was not detected in three genomes (Cloacimonetes bacterium SCGC AAA252-P10, Candidatus Cloacimonas sp. 4484-209, and CLC_T1 Bin 23_1). Pyruvate is predicted to be converted to acetyl-CoA by pyruvate ferredoxin oxidoreductase (PFOR) (EC: 1.2.7.1), which was encoded in 98% of the genomes (45/46). Additionally, 48% (22/46) of the Cloacimonadota genomes encode a predicted pyruvate dehydrogenase (EC: 1.2.4.1), underscoring the likely importance of acetyl-CoA generation within Cloacimonadota metabolism.

The Cloacimonadota genomes encode highly incomplete tricarboxylic acid (TCA) cycles, with only two of the genes in the TCA cycle found in more than 22% of the genomes. Fumarate hydratase (EC: 4.2.1.2), which catalyzes the conversion of fumarate to malate, was found in 89% of the genomes (41/46), and oxoglutarate ferredoxin oxidoreductase (EC: 1.2.7.3 or 1.2.7.11), catalyzing the formation of succinyl-CoA from 2-oxoglutarate, was identified in 96% of the genomes (44/45). The Cloacimonadota genomes typically possess part of the pentose phosphate pathway, particularly the genes transketolase (EC: 2.2.1.1), ribulose-phosphate 3-epimerase (EC: 5.1.3.1), ribose 5-phosphate isomerase A (EC: 5.3.1.6), ribose-phosphate pyrophosphokinase (EC: 2.7.6.1), and phosphoglucomutase (EC: 5.4.2.2), with each being present in 80 to 96% of genomes (37 to 44/46) highlighting a capacity for sugar transformation.

Sugars may derive from more complex carbohydrates - Cloacimonadota genomes encode a suite of carbohydrate-active enzymes (CAZy), the enzymes involved in cleavage of specific sugar linkages (Lombard *et al*., 2014). Prominent CAZy enzymes encoded in Cloacimonadota genomes include polyphenolics, starch, and chitin cleavage proteins from the DRAM annotation (Figure S1). The prevalence of these CAZy enzymes was higher in the genomes from Clade I, suggesting Cloacimonadota in engineered environments can each individually catabolize a wider variety of substrates than their Clade II counterparts, perhaps reflective of the higher heterogeneity of the available substrates in these engineered systems. To further investigate CAZy enzymes the genomes were annotated with dbCAN2 (Zhang *et al*., 2018). The CAZy modules with the highest number of genes in Cloacimonadota genomes were glycosyltransferases and glycoside hydrolases (Figure S2). Unique to Clade I are auxiliary activities (Figure 4, pink). Specifically, CAZy family AA6 was encoded in 69% (24/35) of Clade I genomes and is classified as 1,4-benzoquinone reductase (EC: 1.6.5.6) in the CAZy database, where in *Burkholderia* sp. the enzyme plays a role in 4-aminophenol metabolism (Takenaka *et al*., 2011).

**Figure 4.**
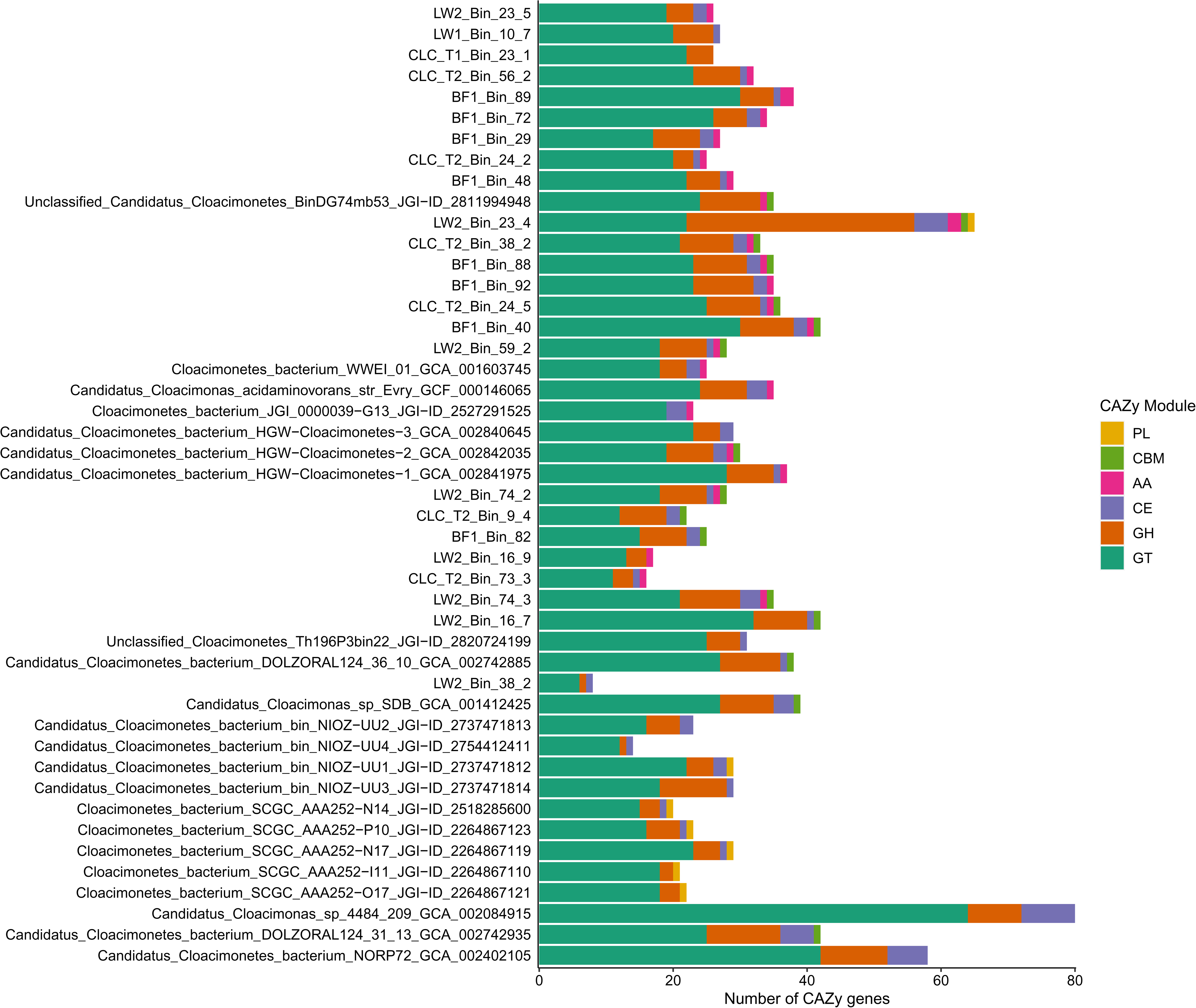
Barplot depicting count of carbohydrate-active genes identified from each Cloacimonadota genome, based on dbCAN gene annotations from the CAZy database. Genomes are ordered as in the phylogeny in Figure 2. CAZy modules are colored to distinguish different functions. PL (yellow) = polysaccharide lyases; CBM (green) = carbohydrate binding modules; AA (pink) = auxiliary activity; CE (purple) = carbohydrate esterases; GH (orange) = glycoside hydrolases; GT (teal) = glycosyltransferases.

Another important aspect of Cloacimonadota carbon metabolism involves acetate. The acetogenic pathway requires phosphate acetyltransferase and acetate kinase, with concomitant substrate-level phosphorylation of ADP to ATP. Phosphate acetyltransferase (EC: 2.3.1.8) was present in 91% of genomes (42/46). Acetate kinase (EC: 2.7.2.1) was present in 72% of all Cloacimonadota genomes (33/46). However, the presence of acetate kinase is not evenly distributed across the phylogenetic tree, where 91% of the Clade I genomes (32/35) encode acetate kinase while this gene was not detected in the Clade II genomes. Acetate kinase was detected in one of the three more basal genomes outside Clade I and Clade II (Candidatus Cloacimonetes bacterium DOLZORAL_124_31_13). Some Cloacimonadota genomes encode the potential to regenerate acetyl-CoA through acetyl-CoA synthetase (EC: 6.2.1.1), which was present in 22% of genomes (10/46). Our analyses suggest that Cloacimonadota are not capable of generating acetyl-CoA via the Wood Ljundahl pathway, as the CO-methylating acetyl-CoA synthetase (EC: 2.3.1.169) was not detected in any genome. Intermediate steps of the Wood-Ljungdahl pathway, methenyltetrahydrofolate cyclohydrolase (EC: 3.5.4.9) and methylenetetrahydrofolate dehydrogenase (EC: 1.5.1.5), were detected in all Cloacimonadota genomes, and formate-tetrahydrofolate ligase (EC: 6.3.4.3) was detected in 61% of genomes (28/46). Though these genes can be associated with carbon fixation in conjunction with CO-methylating acetyl-CoA synthetase, we expect these genes are indicative of folate uptake for DNA synthesis in these organisms given dihydrofolate reductase (EC: 1.5.1.3) was detected in 65% of genomes (30/46).

*Ca.* S. thermopropionivorans was identified within a propionate-degrading, methanogenic enrichment culture, where it may play an important role in anaerobic mineralization of organic matter (Dyksma and Gallert, 2019). The genome of *Ca.* Syntrophosphaera thermopropionivorans encodes genes for propionate oxidation via methylmalonyl-CoA (Dyksma and Gallert, 2019), though the capacity for propionate oxidation in this organism has not been directly determined. Several genes from this pathway were not detected in *Ca.* S. thermopropionivorans’ genome, including the critical gene propionate-CoA transferase (EC: 2.8.3.1), which converts propionate to propionyl-CoA, and succinyl-CoA synthase (EC: 6.2.1.5). Similarly, these genes were detected in 0 and only 17% of Cloacimonadota genomes (8/46), respectively. Other important genes in this pathway were prevalent in Cloacimonadota genomes, such as propionyl-CoA carboxylase (EC: 6.4.1.3; 89% of genomes), methyl malonyl-CoA/ethyl malonyl-CoA epimerase (EC: 5.1.99.1; 78%), and methylmalonyl-CoA mutase (EC: 5.4.99.2; 93%). Some enzymes in this pathway were present more patchily, such as succinate dehydrogenase (EC: 1.3.5.1; 22%) and malate dehydrogenase (EC: 1.1.1.37; 9%). Pyruvate carboxylase (EC: 6.4.1.1) was encoded in 96% of genomes (44/46), and pyruvate ferredoxin oxidoreductase (PFOR; EC: 1.2.7.1), as mentioned above, was encoded in 98% of genomes (45/46). Collectively, Cloacimonadota genomes encoded a fragmented pathway for propionate oxidation.

Overall, most Cloacimonadota genomes encode the capacity for sugar cleavage from more complex carbohydrates, sugar fermentation to acetate, and encode a fragmented set of genes involved in the methylmalonyl-CoA pathway for propionate oxidation.

### Energy metabolism

Cloacimonadota are likely fermenters, though for five genomes (11%), fermentation is not the sole predicted mechanism for energy generation. All of the genomes surveyed here lack complexes from the oxidative electron transport chain, including absence of a canonical Complex I (NADH dehydrogenase, EC: 7.1.1.2). None of the genomes encoded a full 14-subunit NADH-quinone oxidoreductase (NuoA, B, C, D, E, F, G, H, I, J, K, L M, and N). One genome, CLC_T2 Bin 9_4, did not encode any subunits, while all other genomes encoded at least one subunit. The most complete NADH dehydrogenases were in the genomes *Ca.* Cloacimonetes bacterium HGW Cloacimonetes-3 (encoding nuoBCDEFGHL), and *Ca.* Cloacimonetes bacterium NORP72 (encoding nuoBCDHILMN), each with eight subunits (57%). The former, *Ca.* Cloacimonetes bacterium HGW Cloacimonetes-3, contains the NuoEFG module and therefore has the potential for NADH oxidation (Deusch *et al*., 2019). Complex II, succinate dehydrogenase (EC: 1.3.5.1), was detected in 88% of genomes in the environmental Clade II (7/8) but was found in only one of the genomes in the engineered Clade I (*Ca.* Cloacimonetes NIOZ-UU2). Complex III, the cytochrome *bc*_1_ complex (EC: 7.1.1.8), was absent from all genomes. Complex IV, cytochrome *c* oxidase (EC: 7.1.1.9), was absent in all genomes except for *Ca*. Cloacimonetes bacterium NORP72. Complex V, ATP synthase, occurs in two forms. The F- type ATPase (alpha, beta, gamma, delta, epsilon, a, b, c – 8 subunits) was partially present in 23% (8/35) of the Clade I genomes. In contrast, 88% (7/8) of the environmental Clade II genomes contained at least partial complexes, with five of eight genomes encoding complete F-type ATPases (Figure S1). The V/A-type ATPase (A, B, C, D, E, F, G/H, I, K – 9 subunits), was partially detected in 86% (30/35) of the engineered Clade I genomes and was completely absent in the Clade II genomes, though notably, no complete V/A-type ATPases were detected in any Cloacimonadota genome (Figure S1). Clade I genomes encoding a V/A-type ATPase is consistent with earlier metabolic predictions from Cloacimonadota genomes from engineered ecosystems (Dyksma and Gallert, 2019).

ATP synthases rely on generation of a membrane potential, and the absence of a canonical electron transport chain indicates an alternate mechanism is used by Cloacimonadota. The Rnf electron transport complex is a ferredoxin:NAD oxidoreductase that oxidizes reduced ferredoxin and reduces NAD, generating a transmembrane ion gradient, in a reversible reaction (Hess *et al*., 2016). Each of the genes from the Rnf complex (*rnf*ABCDEG) were encoded in 85 to 93% (39 to 43/46) of Cloacimonadota genomes, with 70% (32/46) of genomes encoding the complete complex. The Rnf complex can be important for bacteria in terms of energy conservation, and along with an ATP synthase, creates a simple two-step respiratory chain in the fermentative *Thermotoga maritima* (Kuhns *et al*., 2020). Two genomes, LW2 Bin 23_5 and *Ca.* Cloacimonetes bacterium NORP72, did not encode any components of the complex. The NORP72 genome contained a more-complete canonical electron transport chain than any other Cloacimonadota, suggesting this basal lineage may generate a membrane potential through NADH dehydrogenase and cytochrome C oxidase. For LW2 Bin 23_5, this may indicate an absence due to incomplete assembly or binning, as it is predicted to be 84% complete.

Hydrogenases are an alternative mechanism for generating a membrane potential. Ferredoxin hydrogenase (EC: 1.12.7.2) was present in 89% of the Clade I Cloacimonadota genomes (31/35), and none of the Clade II genomes. Of the three genomes that did not fall into either clade, a ferredoxin hydrogenase was detected only in *Ca*. Cloacimonetes bacterium DOLZORAL_124_31_13. There was no overlap between the presence of ferredoxin hydrogenases and complete ATP synthases. Additionally, formate dehydrogenase (EC: 1.17.1.9) was encoded in 74% of Clade I Cloacimonadota genomes (26/35), one Clade II genome (Cloacimonetes bacterium SCGC AAA252-N17), and one genome not in either clade (*Ca*. Cloacimonetes sp. 4484_209). Other [NiFe] and [FeFe] hydrogenases (K00437 and K17997, respectively) were not detected in any genomes. The functionality of complex V in the presence of membrane-potential-generating mechanisms is possible due to the presence of the Rnf complex, but the significance of hydrogenases in Cloacimonadota energy generation is unclear due to their inconsistent distribution across genomes and clades.

Cloacimonadota genomes largely lack any genes involved in nitrogen, sulfur and methane transformations for energy generation. For nitrogen cycling, 80% of the engineered Clade I genomes (28/35) have predicted hydroxylamine reductases (EC: 1.7.99.1), or ammonia:acceptor oxidoreductases, which were absent in all Clade II genomes. The majority of genomes lack genes involved in denitrification or dissimilative reduction of nitrate. The only exception was the genome *Ca.* Cloacimonetes bacterium NORP72, which encodes both a predicted nitrate reductase and a predicted nitric oxide reductase – these may connect to the encoded partial electron transport chain, allowing this organism, uniquely among the Cloacimonadota, to respire using nitrate as a terminal electron acceptor. Cloacimonadota genomes generally lacked genes involved in sulfur metabolism and sulfur cycling, but the environmental Clade II genomes did show patchy presence of some relevant genes. Specifically, 88% (7/8) of genomes had both an anaerobic sulfite reductase (*asr*) and a sulfur reductase (EC: 1.12.98.4). The Clade II genome that lacked both of these genes was Cloacimonetes bacterium SCGC AAA252-P10. These predicted functions point to a role for Cloacimonadota from marine and anoxic lake environments in hydrogen sulfide production. All Cloacimonadota genomes lacked haloacid dehalogenases (E.C: 3.8.1.2), reductive dehalogenases (EC: 1.21.99.5), and arsenic reductases (EC: 1.20.4.1), and only 17% of genomes (8/47) contain predicted selenate reductases (EC:1.97.1.9).

A previous study suggested that Cloacimonadota ferment amino acids to generate energy, pointing to five ferredoxin oxidoreductases as key markers for this activity (Pelletier *et al*., 2008). The five enzymes include: pyruvate ferredoxin oxidoreductase (K03737), 2-ketoglutarate ferredoxin oxidoreductase (K00174, K00175, K00176, and K00177), aldehyde ferredoxin oxidoreductase (K03738), branched-chain ketoacid ferredoxin oxidoreductase (K00186, K00187, K00188, and K00189), and indolepyruvate ferredoxin oxidoreductase (K00179, and K00180). All five ferredoxin oxidoreductases were encoded in at least some Cloacimonadota genomes, though with varying distributions across clades. Abundant ferredoxin oxidoreductases in all genomes included pyruvate ferredoxin oxidoreductase (K03737, 98% of genomes (45/46)) and indolepyruvate ferredoxin oxidoreductase (K00179, 91% of genomes (42/46)). Three subunits of 2-ketoglutarate ferredoxin oxidoreductase (*kor*A, *kor*B and *kor*D) were each encoded in over 93% of genomes, but one subunit (*kor*C) was only encoded in 24% of genomes (11/46). Aldehyde ferredoxin oxidoreductase was encoded in 66% of Clade I genomes (23/35) but none of the Clade II genomes. In contrast, branched chain ketoacid ferredoxin oxidoreductase (*vor*) was not detected in Clade I genomes and was patchily present in Clade II genomes. Two of the four subunits, *vor*A and *vor*C, were both encoded in 88% of Clade II environmental genomes (7/8), while the other two subunits, *vor*B and *vor*G, were not encoded in any genome. Overall, the presence of ferredoxin oxidoreductases suggests the capability of amino acid fermentation encoded within Cloacimonadota genomes, with some variation between clades. To evaluate the distribution of peptidases and peptidase inhibitors, BLASTP was used to search each of the genomes against the MEROPS database (Rawlings *et al*., 2018). The peptidase families with the highest number of genes in Cloacimonadota genomes were the metallo, serine and cysteine peptidases (Figure S3). Cloacimonadota genomes encode a variety of peptidases with no strong delineation between clades.

From the genomic assessments of metabolic capacity, we predict that these Cloacimonadota are obligate anaerobic acetogenic organisms, without major roles in the sulfur and nitrogen cycles (Figure 5). The examined Cloacimonadota genomes contain the genes PFOR and acetate kinase, involved in conversion of pyruvate to acetate. However, acetate kinase was only prevalent in Clade I, suggesting differences in energy generation between the two clades defined by the phylogenetic tree. For energy production via ATPases, the environmental Clade II organisms encode the F-type ATPase, whereas the engineered Clade I organisms lack a complete ATPase but have partial V/A type ATPases. Succinate dehydrogenase was frequently identified in Clade II genomes, while formate dehydrogenase was consistently detected in Clade I genomes, indicating different mechanisms of generating a membrane potential. Most Cloacimonadota encoded the RNF complex, which also may help create a transmembrane proton gradient, influencing energy generation. The presence of ferredoxin oxidoreductases and a variety of peptidases suggest Cloacimonadota may play a role in the degradation of proteins and/or amino acids.

**Figure 5.**
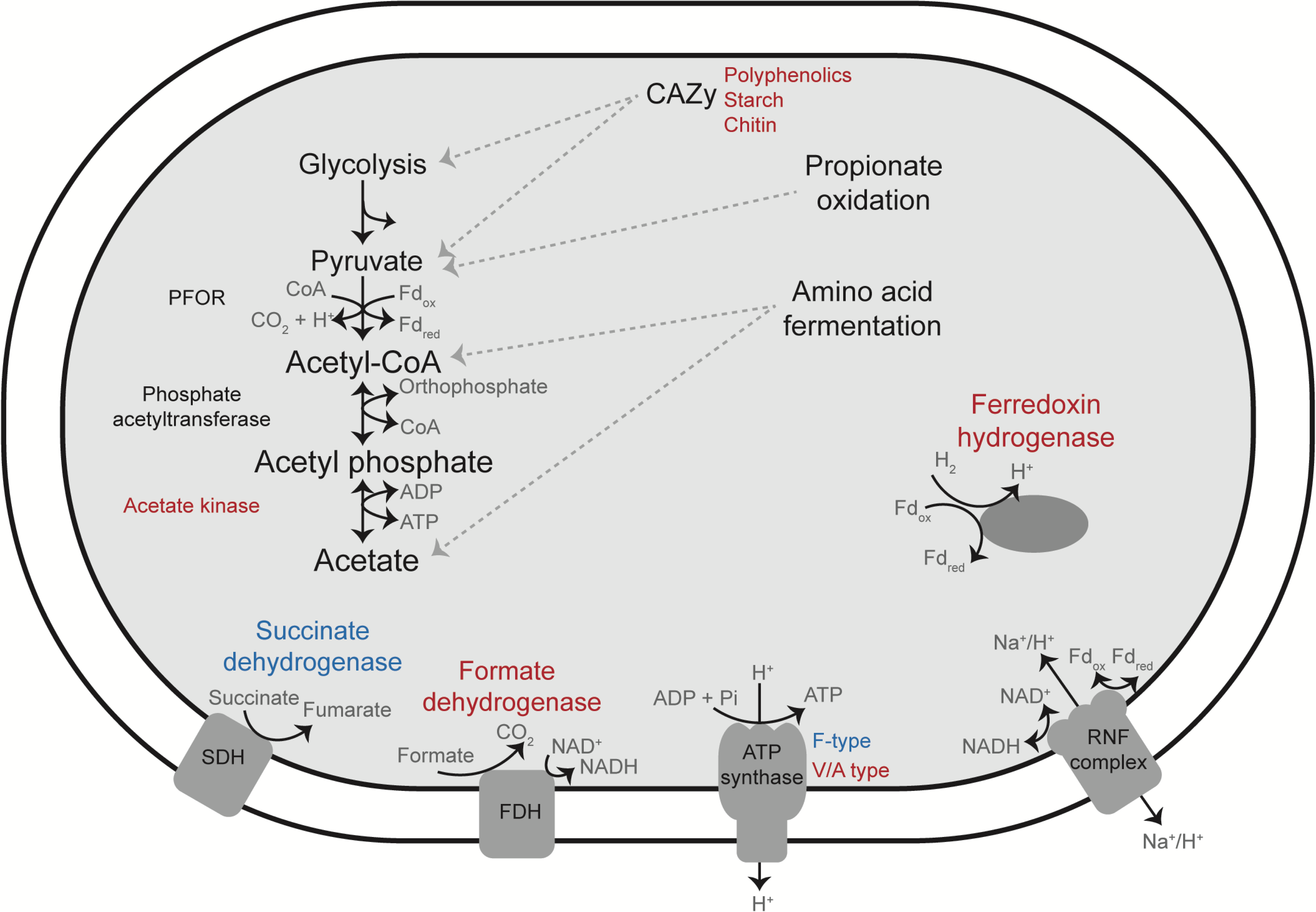
Simplified cell diagram of carbon and energy metabolism in Cloacimonadota reconstructed from annotations of assembled MAGs and reference genomes. Red labels indicated the gene was encoded prominently in Clade I (largely engineered) genomes, and blue labels indicated genes which were encoded prominently in Clade II (environmental) genomes. Black labels specify proteins encoded in all Cloacimonadota genomes. Dotted lines denote possible paths for substrate degradation. Not all genes were detected in the glycolysis or propionate oxidation pathways (see text for details). Amino acid degradation is indicated by the presence of ferredoxin oxidoreductases. CAZy = carbohydrate-active enzymes; PFOR = pyruvate ferredoxin oxidoreductase; SDH = succinate dehydrogenase; FDH = formate dehydrogenase.

### Cloacimonadota pangenome

The phylogenetic separation of engineered and environmental ecosystem-derived organisms was unexpected, so we examined the pangenome of the phylum to assess traits that separated these groups and might explain adaptations specific to each habitat type.

The anvi’o analysis platform was used to create a pangenome for the 46 high quality genomes included in the metabolic analyses (Figure 6). The pangenome analysis used pairwise amino acid similarity searches to identify gene clusters (Delmont and Eren, 2018). Our pangenome analysis resulted in 5,149 gene clusters from a total of 64,300 genes across the 46 genomes, capturing 70% of genes in a gene cluster. Each of the gene clusters contain genes from a minimum of three genomes, as singletons and doubletons were excluded from this pangenome analysis (parameter --min_occurrence = 3). To highlight patterns of shared genomic profiles, genomes were ordered based on the frequency of the gene clusters that they share (Figure 6). The genomes largely clustered by the ecosystem type from which the genome originated, similar to the phylogenetic tree (Figure 2), with some exceptions discussed below. We manually selected groupings of gene clusters based on highly conserved gene cluster patterns across the Cloacimonadota genomes (Groups 1-4, Figure 6).

**Figure 6.**
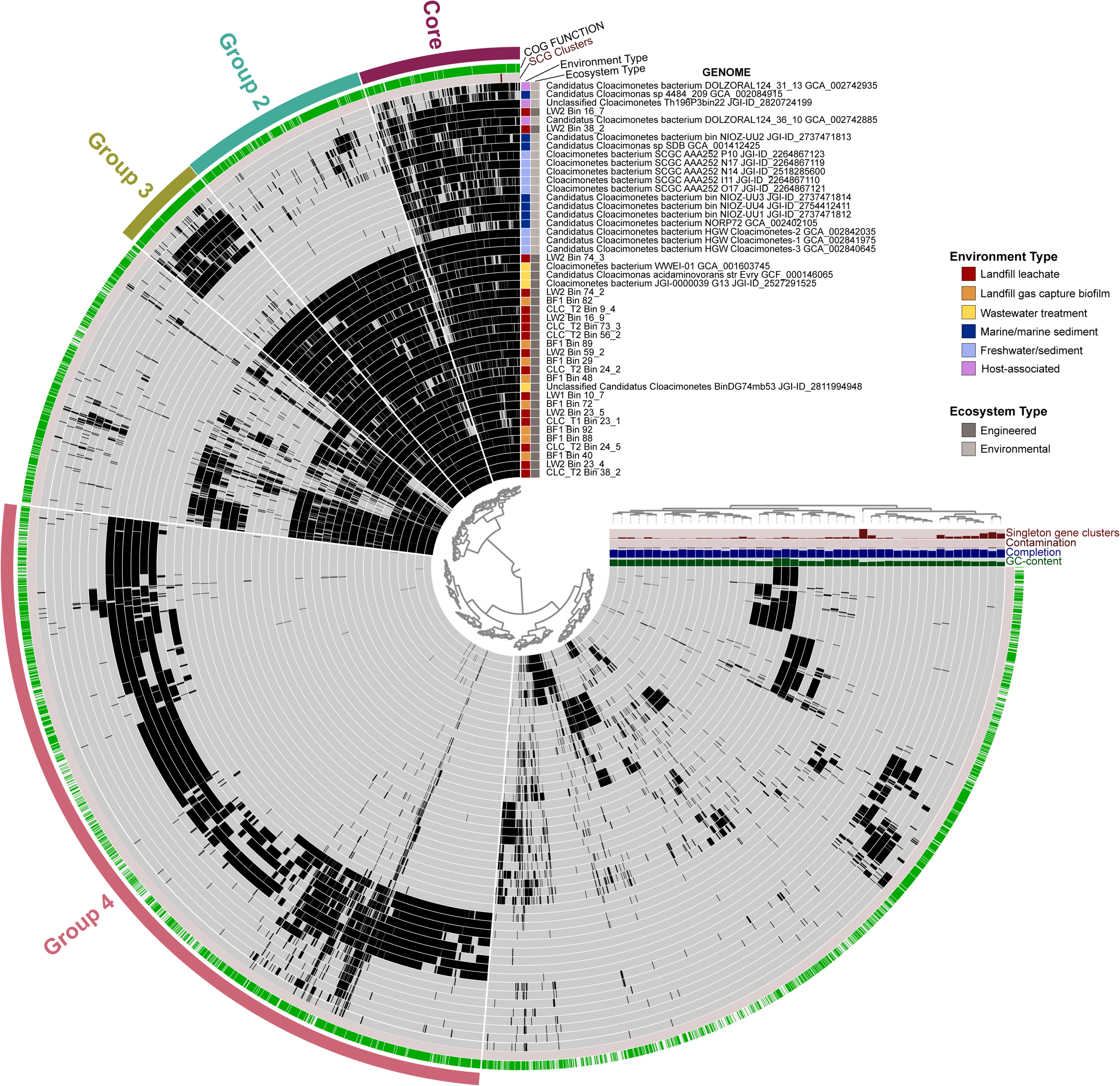
Cloacimonadota pangenome with 5,149 gene clusters from 64,300 genes identified from 46 genomes. Genomes are organized based on the frequency of the gene clusters that they share, with each genome represented as a ring on the plot and genes marked as present (black) or absent (grey) within gene clusters (rays around the circle). The gene clusters represent a minimum of three genomes, as singletons and doubletons were excluded from the visualization. The ‘core’ wedge (Group 1) corresponds to the 347 gene clusters that were near-universal in genomes originating from engineered ecosystems. The “Group 2” wedge corresponds to 405 clusters that were nearly universal in genomes originating from engineered ecosystems. The “Group 3” wedge corresponds to 202 gene clusters that are nearly universal in genomes from engineered ecosystems and six other genomes that are phylogenetically associated with Clade I on the phylogenetic tree. The “Group 4” wedge is a collection of 1,765 gene clusters from genomes that originated from an environmental ecosystem. Bar graphs on the right indicate parameters for each genome: GC content (%), completion (%), contamination (%), and number of singleton gene clusters. Colored boxes to the left of the genome names indicate the environment type and ecosystem type from which the genome originated.

Each gene cluster group was examined based on the COG categories for the represented gene clusters (Figure S4). Group 1, labelled ‘core’, contains gene clusters that are near-universal and highly conserved across all Cloacimonadota, and contains 347 gene clusters. The “core” section contained the highest number of COG category “J – translation” genes. Other categories of essential functions were also well represented, such as “C -energy production and conversion”, “F – nucleotide metabolism and transport”, “L – replication and repair”, “M – cell wall/membrane/envelope biogenesis”, and “O – post-translational modification”. Interestingly, “R – general function prediction only” encompasses 6% of the “core” wedge, identifying highly conserved genes with no known function that are likely important for Cloacimonadota growth or survival.

Group 2 contained 405 gene clusters that were nearly universal in genomes originating from engineered ecosystems but largely absent in genomes from other environments. This section had a comparatively high proportion of genes in “M – cell wall/membrane/envelope biogenesis” and “P – inorganic ion transport” (Figure S4). The Group 2 gene clusters included a copper chaperone (*cop*Z) and a copper oxidase (laccase) domain. Copper chaperones are responsible for transportation and transfer of copper (Rosenzweig and O’Halloran, 2000), and laccase is a copper oxidase that catalyzes oxidation of a broad range of compounds (Reiss *et al*., 2013). Engineered environments typically have higher metal concentrations, which may select for metal tolerance mechanisms. Another interesting gene cluster was chromate transport protein (*chr*A). Chromate transport is an important microbial heavy metal resistance mechanism for managing chromium (Díaz-Pérez *et al*., 2007), and may be of importance to Cloacimonadota in engineered environments. Other gene clusters, such as ABC-type Fe^3+^ transporters and divalent metal cation (Fe/Co/Zn/Cd) transporter are present in both the ‘Engineered’ and ‘Environmental’ sections (see below), indicating these functions are important across the Cloacimonadota, but had diverged sufficiently to form separate gene clusters within the pangenome analysis.

Group 3 contained 202 gene clusters that were nearly universal in genomes that correspond to Clade I on the phylogenetic tree. The pangenome Group 3 gene clusters have representation from genomes from engineered ecosystems (interior rings) as well as an additional six genomes (exterior rings) (Figure 6), which all correspond to Clade I (Figure 2). The COG category that is comparatively abundant in this group was “J – translation”, similar to the ‘core’ Group 1. It is possible for functional overlap across the groups or with gene clusters outside of the defined groups, due to sequence dissimilarity that drives the generation of separate gene clusters. In the case of Group 3, we did not identify any key functions that were specifically associated with the genomes represented in the group – this group appears to indicate higher sequence conservation for these functions among the Clade I organisms rather than specific gains or losses of function.

The last identified grouping, Group 4, contains 1,765 gene clusters from genomes originating from environmental ecosystems, and is thus the largest identified Group. A notable category in Group 4 is “V - defense mechanisms”. The ‘core’, Group 2, and Group 3 sections each contained a small number of genes, including ABC-type antimicrobial transport systems and a multidrug efflux pump. In contrast, Group 4 contained gene clusters for components of the CRISPR/Cas system. However, CRISPR/Cas system genes were also detected in genomes originating from engineered environments and the landfill MAGs. The presence of a CRISPR/Cas system in Cloacimonadota genomes does not appear to be clearly linked to the functional or phylogenetic distribution of Cloacimonadota, but the CRISPR/Cas system does appear more tightly conserved in the environmentally-derived genomes, allowing for identification of gene clusters from Clade II genomes.

The presence of the same genes across multiple gene clusters is due to sequence dissimilarity breaking gene clusters into two or more clusters with higher sequence identities. We were evaluating a phylum-level lineage, where pangenome comparisons are better targeted to species- or strain-level comparisons (Delmont and Eren, 2018). However, the high number of identified gene clusters, and the connections between ecosystem and gene complements do provide interesting insights into Cloacimonadota evolution and specialization. Notably, eight genomes (Figure 6, exterior circles) clustered differently based on gene clusters compared to their placement on the phylogenetic tree (Figure 2). First, the six genomes *Ca.* Cloacimonas sp. SDB, *Ca.* Cloacimonetes bacterium NIOZ-UU2, *Ca.* Cloacimonetes bacterium DOLZORAL_124_36_10, Unclassified Cloacimonetes Th196P3bin22, along with landfill MAGs LW2 Bin 38_2 and LW2 Bin 16_7, clustered together due to a unique pattern of shared gene clusters, with strong representation in Group 3 (gene clusters shared by engineered ecosystem-derived genomes), a partial presence in Group 4 (gene clusters from environmentally-derived genomes), and a third, discrete cluster of genes that is more patchily represented across the six genomes (Figure 6, 4 o’clock). Notably, this set of genomes contains two of the three host-associated Cloacimonadota, as well as two landfill-derived MAGs and two marine-associated MAGs, suggesting there may be a shared host-associated lifestyle for these organisms. All of these genomes were affiliated with Clade I, engineered environments, on the phylogenetic tree. Further, the three host-associated genomes (*Ca.* Cloacimonetes bacterium DOLZORAL_124_31_13, *Ca.* Cloacimonetes bacterium DOLZORAL_124_36_10, Unclassified Cloacimonetes Th196P3bin22) along with landfill MAG LW2 Bin 16_7 and *Ca.* Cloacimonetes bacterium NORP7, were five genomes that encoded many genes involved in a flagellar system as determined through the KEGG annotation (Table S2). Flagella may be a trait of host-associated Cloacimonadota which is contributing to the organization of genomes in the pangenome.

Three genomes have the lowest number of overall shared gene clusters and are not affiliated with any group. These three genomes are also the three genomes that fall outside Clade I and Clade II, placing as basal and most divergent on the phylogenetic tree (Figure 2). Of these, *Ca.* Cloacimonas sp. 4484_209 and *Ca.* Cloacimonetes bacterium DOLZORAL_124_31_13 (furthest exterior rings, Figure 6), seem to be outliers because they share ‘core’ gene clusters but lack gene clusters from other major groupings. In contrast, *Ca.* Cloacimonetes bacterium NORP72 (middle of genome rings, Figure 6), does not share many gene clusters with the other genomes and contains the largest number of singleton genes. *Ca.* Cloacimonetes bacterium NORP72 also contains fewer hits across all the COG categories for Groups 1-4 compared to the other genomes (Figure S4). *Ca.* Cloacimonetes bacterium NORP72 was the only genome to encode certain genes, such as nitrate reductase, and was a genome with the largest genome, largest number of genes, and longest branch lengths to other Cloacimonadota genomes (Figure 2). Collectively these differences suggest *Ca.* Cloacimonetes bacterium NORP72 is either a representative of a distantly related Cloacimonadota clade (Class level difference), or possibly belongs within a different phylum.

## Conclusions

Here we reconstructed 24 Cloacimonadota MAGs from a landfill site, expanding the number of Cloacimonadota genomes by 109% and more than doubling the number of good quality genomes available. Cloacimonadota MAGs were assembled from landfill leachate wells, a composite leachate cistern, and a biofilm sample from the landfill site, often at low proportional abundances. The MAG with the highest proportional abundance was obtained from the biofilm sample, where it accounted for 4.72% of the assembled metagenome (BF1 Bin 72). In a phylum-level comparison of phylogenetic diversity, Cloacimonadota genomes formed two distinct clades segregated by ecosystem, with one clade containing all engineered-environment derived genomes and one clade containing the majority of environmentally derived genomes. From metabolic predictions, Cloacimonadota are predicted to be anaerobic, acetogenic, fermentative bacteria. Substrate utilization may differ across lineages; for example, carbohydrate cleavage enzymes were more diverse in genomes from engineered environments, suggesting different substrate capacities. Propionate oxidation via methylmalonyl-CoA may be an important function based on culture trials, but the pathway was only partially encoded in most genomes. Ferredoxin oxidoreductases suggest the capacity for amino acid fermentation. Most Cloacimonadota genomes encode a RNF complex, which may be important in energy generation. ATPase type was phylogenetically linked, where Clade I genomes lacked complete ATPase but had a partial V/A-type ATPase, and some genomes from Clade II encoded a F-type ATPase. This indicates there may be clade specific mechanisms for energy generation. Cloacimonadota genomes from engineered environments were enriched in metal transport and resistance genes and share substantially higher sequence identities compared to their environmentally derived counterparts. Pangenome and phylogenetic analyses indicate a shared evolutionary history for Cloacimonadota from engineered environments, where this lineage is frequently identified and often at high abundance.

## Experimental procedures

### Sampling, DNA sequencing, assembly and binning

Samples were collected from an active municipal landfill site in Southern Ontario. Samples included landfill leachate wells (n = 5), groundwater wells (n = 1), and a biofilm (n = 1) collected over three separate sampling events. In the first sampling in July 2016, leachate biomass was collected from a composite leachate cistern (CLC_T1) by filtration of the leachate through a 0.2 µm and a 0.1 µm polyethersulfone filter in series. One week later, leachate biomass was collected from the same composite leachate cistern (CLC_T2) along with three leachate wells (LW1, LW2, and LW3), and one groundwater well (GW1) by filtration through a 3 µm glass fiber filter followed by a 0.1 µm polyethersulfone filter in series. This change in filter pore sizes was made to reduce filter clogging. Glass fiber pre-filters were discarded, and 0.1 µm filters were frozen on dry ice and transported to the lab where they were stored at -80°C until processed. A biofilm sample (BF1) was collected from overgrowth in the landfill’s gas capture system in October 2017. The sample was extricated from a clogged valve using sterile scalpels, as is resisted any other form of manipulation.

DNA was extracted from the landfill biomass on filters and the gas intake biofilm sample using a Qiagen PowerSoil DNA isolation kit following the manufacturer’s instructions, with the modification of slicing the filters into pieces and adding the filter directly to the extraction in place of soil, or with the addition of 10 g of biofilm in place of soil. DNA was shotgun sequenced on a Illumina HiSeq machine with 150 bp paired-end reads (U.S. Department of Energy Joint Genome Institute).

For all samples collected in 2016, the metagenomic sequences were assembled with IDBA_UD (Peng *et al*., 2012), and binned with CONCOCT (Alneberg *et al*., 2014) using tetranucleotide frequencies and cross-sample abundance patterns based on read mapping. Bins were manually refined on the anvi’o analysis platform (Eren *et al*., 2015). The biofilm metagenome from 2017 was assembled with metaSPAdes (Nurk *et al*., 2017), and binned using DasTool (Sieber *et al*., 2018) taking input from the binning algorithms CONCOCT, MaxBin2 (Wu *et al*., 2016), and MetaBAT2 (Kang *et al*., 2019). All MAGs were required to have thresholds of completion above 70% and contamination below 10% to be considered for metabolic analyses. MAGs were identified as belonging to the members of the phylum Cloacimonadota based on placement within a concatenated ribosomal protein tree that contained representatives across the tree of life (Sauk and Hug, 2021). The bin average fold coverage was calculated for each MAG based on read mapping of the input reads to the relevant metagenome using Bowtie2 v. 2.3.5.1 (Langmead and Salzberg, 2012).

### Mining public databases for Cloacimonadota genomes

All publicly available genomes for members of the phylum Cloacimonadota were downloaded from the Joint Genome Institute’s Integrated Microbial Genomes and Microbiomes (http://img.jgi.doe.gov) or the National Center for Biotechnology Information (http://www.ncbi.nlm.nih.gov) (July 2019). Three genomes for members of the Gemmatimonadota, five genomes for members of the Fibrobacterota and six genomes for members of the Zixibacteria were also downloaded as phylogenetically related yet distinct outgroups for phylogenetic trees. Genome quality was evaluated with CheckM (Parks *et al*., 2015) for all collected genomes, and genomes that did not meet the >70% completion and <10% contamination criteria were not included in metabolic analyses.

### Phylogenetic tree

Landfill Cloacimonadota MAGs and all reference genomes were placed on a concatenated ribosomal protein tree. A suite of 16 conserved, single-copy, non-laterally transferred ribosomal proteins (ribosomal proteins L2, L3, L4, L5, L6, L14, L15, L16, L18, L22, L24, S3, S8, S10, S17, and S19) were extracted from each genome (Hug *et al*., 2013). Each gene set was aligned using MAFFT version 7.402 (Katoh, 2002) and positions with ≥ removed to manually curate the alignments. Curated alignments were concatenated, and any sequences that contained less than 50% of the total number of aligned amino acids were removed. A maximum likelihood phylogeny was then inferred using RaxML version 7.471 with automatic bootstrapping under a LG protein substitution model (Stamatakis, 2006). The phylogenetic tree was visualized using iTOL (http://itol.embl.de) (Letunic and Bork, 2016).

### Metabolic analyses

#### KEGG, DRAM, CAZy, MEROPS

Cloacimonadota genomes, both reference genomes from public databases and our landfill MAGs, meeting the thresholds of completion above 70% and contamination below 10% were included in a survey of phylum Cloacimonadota’s metabolic potential (n = 46). Genomes were evaluated using the KEGG Automatic Annotation Server (KAAS) (http://www.genome.jp/kegg/kaas/) (Moriya *et al*., 2007). KAAS was used with bi-directional best hit information based on sequence similarities to provide KEGG orthology identifiers for genes within each genome. Enzyme Commission (EC) numbers or KO numbers were used to cross reference against metabolic functions of interest to ascertain presence or absence (Table S2).

Distilled and Refined Annotation of Metabolism (DRAM) was used under default parameters and without access to the KEGG database, for annotation of the 46 Cloacimonadota genomes (Shaffer *et al*., 2020) in order to assess metabolic predictions in greater detail.

To search for carbohydrate active enzymes, Cloacimonadota genomes were compared to the CAZy database (Lombard *et al*., 2014) using dbCAN2 (version July 31, 2019) (Zhang *et al*., 2018). Annotation were considered if the coverage was greater than 0.5 and e-value was below 1e^-20^ for a hmmer hit. The MEROPS database (https://www.ebi.ac.uk/merops/; merops_scan.lib) was used to search for peptidases and peptidase inhibitors (Rawlings *et al*., 2018). The database was searched with BLASTP v2.8.1+ and hits were considered if the e-value was below 1e^-10^. Beyond these two databases, manual BLASTP searches were conducted to search for superoxide reductase (WP_133200558.1), rubrerythrin (WP_133200557.1) and a pilin protein (WP_133201237.1) from *Candidatus* Syntrophosphaera thermopropionivorans (Dyksma and Gallert, 2019).

### anvi’o pangenome

We used the pangenome analysis tool from the anvi’o package to examine core genes across the phylum, as well as across Cloacimonadota groups associated with engineered and natural environments (Eren *et al*., 2015; Delmont and Eren, 2018). In anvi’o version 6.2, the 46 Cloacimonadota genomes were used to generate anvi’o genomes storage databases, where open reading frames were identified with Prodigal (Hyatt *et al*., 2010), and genes were annotated using the Clusters of Orthologous Groups (COGs) database (Tatusov, 2000). Next, the ‘anvi-pangenome’ program was used to create the pangenome with NCBI blastp for searching (Altschul *et al*., 1990), a minbit parameter of 0.5, a MCL inflation parameter of 2 (for family-level or higher divergence), and a gene cluster min occurrence parameter of 3 (excluding singleton and doubleton clusters). This step identifies ‘gene clusters’ from the similarity searches of each amino acid sequences in each genome to every other amino acid sequence, and further the occurrence of these gene clusters across genomes. Hierarchical clustering analyses for both gene clusters and genomes used Euclidean distance and Ward clustering. In the pangenome visualization, genomes were organized based on the frequency of the gene clusters that they share. Heatmaps were created from the assigned COG categories for gene clusters for each gene cluster in the pangenome, and were visualized using ggplot2 (Ginestet, 2011).

## Data Availability

Landfill metagenomes are publicly available on the Joint Genome Institute’s Integrated Microbial Genomes database as follows: LW1 (IMG ID: 3300014204), LW2 (3300015214), LW3 (3300014205), CLC_T1 (3300014206), CLC_T2 (3300014203), and Biofilm1 (3300028601).

## Acknowledgements

We would like to acknowledge the time, money, and effort required to generate metagenome assembled genomes, including the work of Dr. Elizabeth Edwards and Dr. Camilla Nesbø (University of Toronto); Dr. Laura Villanueva, Dr. Subhash Yadav, and Dr. Bastiaan von Meijenfeldt (Royal Netherlands Institute for Sea Research), and Dr. Inka Vanwonterghem (University of Queensland), each of whom provided permission for us to work with their Cloacimonadota MAGs.

This work was funded by an NSERC Discovery grant (2016-03686) to LAH. LAJ was supported by a Queen Elizabeth II Graduate Scholarship in Science and Technology, an Ontario Graduate Scholarship, and an NSERC CGS-D award. LAH is supported by a Canada Research Chair.

## Table and figure legends

**Supplementary Figure 1.** Presence or absence of functional pathways encoded in Cloacimonadota genomes generated from the DRAM annotation pipeline. Left-most facets are heat maps depicting carbon metabolism and energy pathways, with presence coded by increasing intensity of blue. Right hand facets depict specific metabolic capacities as presence/absence, with green indicating presence of the trait and blue indicating absence.

**Supplementary Figure 2.** Distribution of genes in each CAZy family per Cloacimonadota genome. CAZy genes were annotated with dbCAN2. The CAZy families are: AA = auxiliary activity; CBM = carbohydrate binding modules; CE = carbohydrate esterases; GH = glycoside hydrolases; GT = glycosyltransferases, and PL = polysaccharide lyases.

**Supplementary Figure 3.** Distribution of peptidases per Cloacimonadota genome based on annotation with the MEROPs database. The MEROPS families are: A = aspartic; C = cysteine; I = inhibitors; M = metallo; N = asparagine; S = serine; T = threonine; and U = unknown.

**Supplementary Figure 4.** COG category distribution per genome for groups identified in the pangenome. Groups correspond to those visually selected from the pangenome and are colored to match Figure 6. Clusters of Orthologous Groups (COGs) were annotated in the anvi’o pipeline. Genomes are organized in the heatmap based on the clustering order determined in the pangenome. COG categories by letter are: **(A)** RNA processing and modification; **(B)** Chromatin structure and dynamics; **(C)** Energy production and conversion; **(D)** Cell cycle control and mitosis; **(E)** Amino acid metabolism and transport; **(F)** Nucleotide metabolism and transport; **(G)** Carbohydrate metabolism and transport; **(H)** Coenzyme metabolism; **(I)** Lipid metabolism; **(J)** Translation; **(K)** Transcription; **(L)** Replication and repair; **(M)** Cell wall/membrane/envelope biogenesis; **(N)** Cell motility; **(O)** Post-translational modification; **(P)** Inorganic ion transport and metabolism; **(Q)** Secondary structure; **(R)** General function prediction only; **(S)** Function unknown; **(T)** Signal transduction; **(U)** Intracellular trafficking; **(V)** Defense mechanisms; **(W)** Extracellular structures; **(X)** Mobliome: prophages, transposons; **(Z)** Cytoskeleton.

**Supplementary Table 1.** Characteristics of Cloacimonadota genomes from the landfill site and public databases (as of July 2019), and reference genomes used in analyses.

**Supplementary Table 2.** Presence or absence of metabolic genes of interest identified in Cloacimonadota genomes from the KEGG annotations.

**Supplementary Table 3.** Annotations for each Cloacimonadota genome obtained with DRAM.

