## Supplemental figures S1-S4 for "Cloacimonadota metabolisms include adaptations for engineered environments that are reflected in the evolutionary history of the phylum"

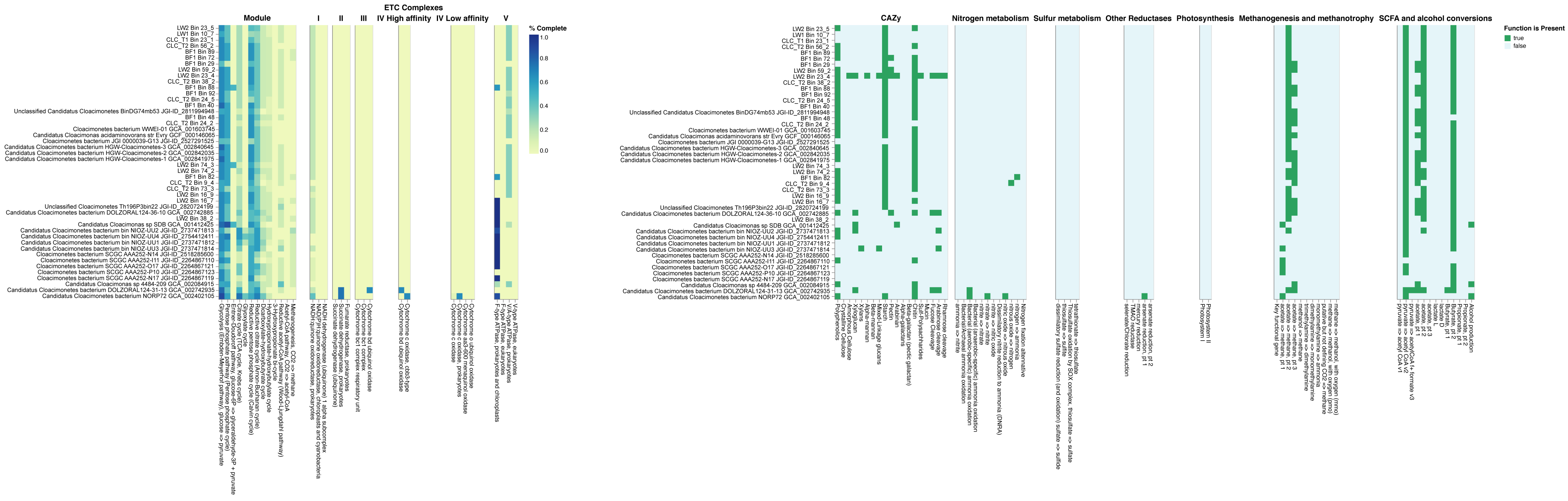


Supplementary Figure 1. Presence or absence of functional pathways encoded in Cloacimonadota genomes generated from the DRAM annotation pipeline. Left-most facets are heat maps depicting carbon metabolism and energy pathways, with presence coded by increasing intensity of blue. Right hand facets depict specific metabolic capacities as presence/absence, with green indicating presence of the trait and blue indicating absence.

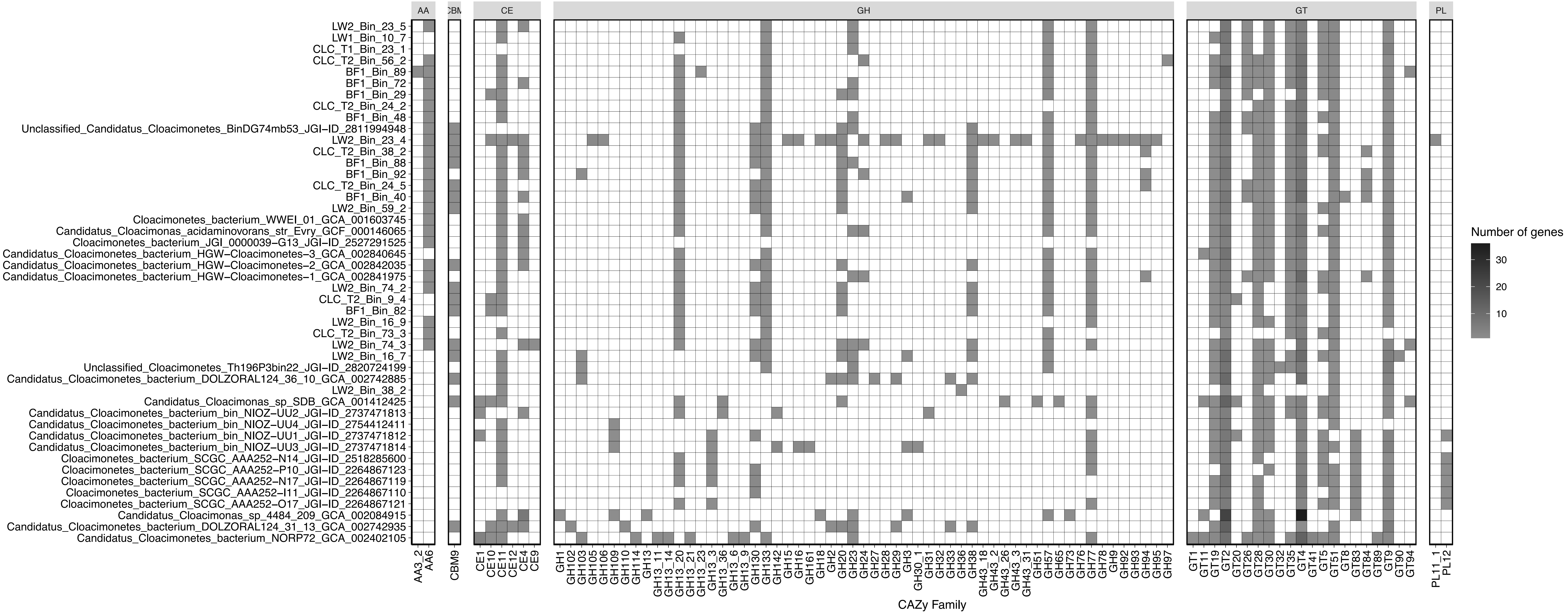


Supplementary Figure 2. Distribution of genes in each CAZy family per Cloacimonadota genome. CAZy genes were annotated with dbCAN2. The CAZy families are: AA = auxiliary activity; CBM = carbohydrate binding modules; CE = carbohydrate esterases; GH = glycoside hydrolases; GT = glycosyltransferases, and PL = polysaccharide lyases.

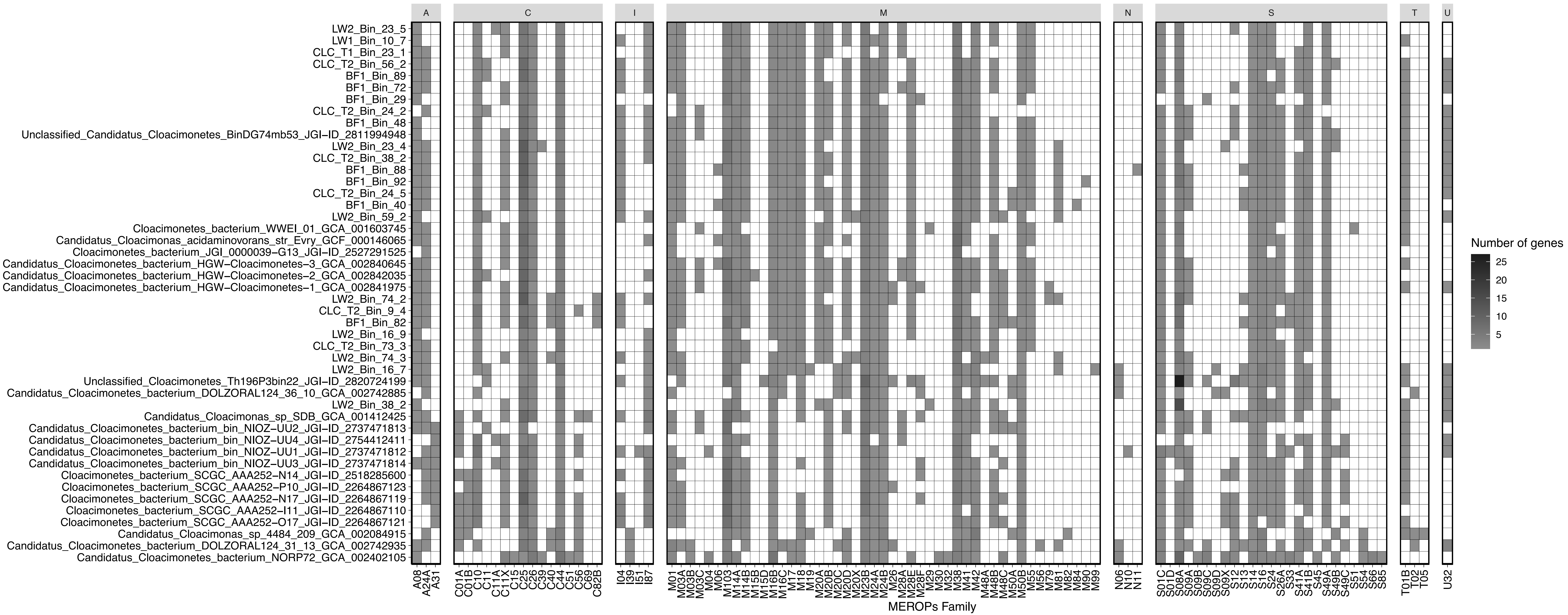


Supplementary Figure 3. Distribution of peptidases per Cloacimonadota genome based on annotation with the MEROPs database. The MEROPS families are: A = aspartic; C = cysteine; I = inhibitors; M = metallo; N = asparagine; S = serine; T = threonine; and U = unknown.

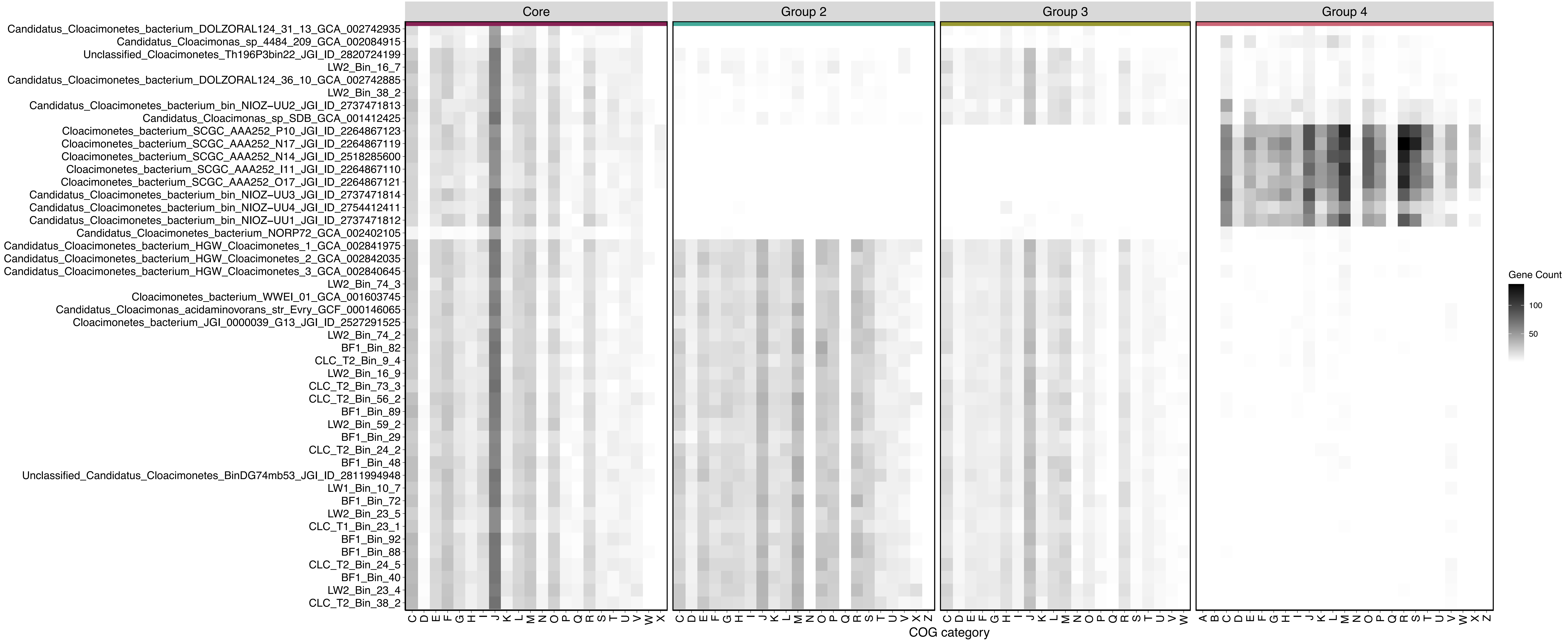


Supplementary Figure 4. COG category distribution per genome for groups identified in the pangenome. Groups correspond to those visually selected from the pangenome and are colored to match Figure 6. Clusters of Orthologous Groups (COGs) were annotated in the anvi'o pipeline. Genomes are organized in the heatmap based on the clustering order determined in the pangenome. COG categories by letter are: (A) RNA processing and modification; (B) Chromatin structure and dynamics; (C) Energy production and conversion; (D) Cell cycle control and mitosis; (E) Amino acid metabolism and transport; (F) Nucleotide metabolism and transport; (G) Carbohydrate metabolism and transport; (H) Coenzyme metabolism; (I) Lipid metabolism; (J) Translation; (K) Transcription; (L) Replication and repair; (M) Cell wall/membrane/envelope biogenesis; (N) Cell motility; (O) Post-translational modification; (P) Inorganic ion transport and metabolism; (Q) Secondary structure; (R) General function prediction only; (S) Function unknown; (T) Signal transduction; (U) Intracellular trafficking; (V) Defense mechanisms; (W) Extracellular structures; (X) Mobliome: prophages, transposons; (Z) Cytoskeleton.
